# Progesterone receptors drive advanced breast cancer phenotypes including circulating tumor-and stem-like cell expansion in the context of *ESR1* mutation

**DOI:** 10.1101/2025.11.20.689560

**Authors:** Thu H. Truong, Noelle E. Gillis, Amy R. Dwyer, Rosemary J. Huggins, Kyla M. Hagen, Sai Harshita Posani, Nuri A. Temiz, Carlos Perez Kerkvliet, Ellie M. Piepgras, Julie H. Ostrander, Geoffrey L. Greene, Carol A. Lange

## Abstract

Endocrine therapy resistance remains a major challenge in the treatment of advanced estrogen receptor positive (ER+) breast cancer. This can be driven by acquired mutations in the estrogen receptor gene (*ESR1*), such as Y537S or D538G, that primarily emerge in patients with prior aromatase inhibitor therapy and results in constitutive estrogen-independent ER activity. Progesterone receptors (PR) are important modifiers of ER activity, in part via direct binding. We previously showed that PR mediates expansion of cancer stem-like cell (CSC) populations and promotes tamoxifen resistance in nuclear ER/PR transcriptional complexes. In this study, we sought to define whether PR function changes in the context of *ESR1* mutations. PR readily interacted with wild type (WT), but not Y537S or D538G ERs. RNA-seq and ChIP-seq studies demonstrated that ER+ breast cancer models expressing Y537S ER exhibited a distinct response to progesterone. CSC populations were enhanced in Y537S ER+ cells compared to WT ER+ cells. PR knockdown demonstrated that this property required PR expression but was unresponsive to antiprogestins. Moreover, we identified PR-dependent transcriptional programs such as the unfolded protein response (UPR) that can be leveraged to target CSC populations in Y537S *ESR1*-mutant breast cancer. The UPR activator ErSO, but not UPR inhibitors, blocked expansion of CSCs in WT as well as Y537S ER*+* models. Together, our findings demonstrate a critical interplay between PR and mutant ER function and provide insight into PR-driven pathways including hyperactivation of the stress-sensing UPR that can be exploited as potential therapeutic avenues in advanced ER+ breast cancer.

## INTRODUCTION

Estrogen receptor positive (ER+) breast cancers account for ∼80% of cases. Standard of care treatment for ER+ breast cancers includes endocrine therapies such as aromatase inhibitors, selective ER modulators (SERMs), and selective ER degraders (SERDs). These therapies work by targeting estrogen (E2) and ER action, and patients respond well initially with a 5-year survival rate of over 89%. However, the benefits of endocrine therapy are not sustained, and patients can develop late recurrence for 10 to 20 years after initial diagnosis (1). *De novo* or acquired resistance can occur in 15-20% or 30-40% of ER+ breast cancer patients respectively, and these patients will experience recurrence and progress to metastatic disease (2). Research has centered on understanding mechanisms of ER escape from endocrine therapies and can be attributed to factors that include altered coregulator interactions (3), crosstalk with growth factor pathways (4), and genomic alterations to ER such as fusions (5) or gain-of-function mutations (6).

About 20% of patients treated with aromatase inhibitors for ER+ breast cancer acquire mutations in the ER ligand binding domain that are present in metastatic tumors (7). The most common ER mutations that occur are tyrosine 537 to serine (Y537S) and aspartic acid 538 to glycine (D538G), which account for nearly 50% of identified mutations (8–10). These two mutations stabilize ER in the E2-like agonist conformation in the absence of hormone, resulting in a constitutively active ligand-independent ER (11). This leads to decreased affinity for E2 as well as ER-targeted treatments and the emergence of endocrine therapy resistance to first and second-line treatments. Understanding the mechanisms of endocrine therapy resistance in the face of hormone deprivation and ultimate independence from estrogen is crucial to overcoming this major clinical problem and improving overall patient outcomes.

About 65-75% of ER+ breast cancers also contain progesterone receptor (PR) positive cells. Total PR levels have historically been used as a clinical biomarker of ER transcriptional activity to predict response to endocrine therapies. Notably, growing evidence now supports the role of PR as an important ER binding partner and a vital modifier of ER activity and subsequent transcriptional gene programs (12,13). PR has two isoforms, PR-A and PR-B, and studies have shown that these individual isoforms have distinct transcriptional activity and biological phenotypes (14,15). PR can be expressed and remain activated even when ER activity is low or blocked with endocrine therapies through E2/ER-independent mechanisms. For example, PRs are phosphorylated by mitogen- or stress-activated protein kinases that are frequently activated in breast cancers. These phosphorylation events subsequently impact PR signaling and transcriptional activity. Further, activated phospho-PR can auto-induce its own expression via the progesterone receptor gene (*PGR*) (16). Previous work from our lab has demonstrated that PR Ser294 phosphorylation promotes expansion and survival of breast cancer stem-like cells (CSCs) (17–19) and drives gene programs associated with endocrine resistance, embryonic stem development, and CSC biology (20). Breast CSCs are a small subpopulation present in recurrent breast tumors (21); these cells are typically non-proliferating or not actively cycling, making them difficult to target and resistant to standard chemotherapy or endocrine therapy. Additionally, ER gain-of-function mutations such as Y537S and D538G express high PR levels in the absence of E2 (22). There are currently no approved PR-targeted therapies to treat ER+ breast cancer despite PR’s emerging role as an independent driver of disease progression (23).

In this study, we sought to define PR action in the context of ER mutations. Our findings indicate that although cells harboring ER mutations have reduced E2 sensitivity, they exhibit a distinct response to progesterone treatment. Moreover, our studies reveal insights into breast CSC populations that are driven by PR in models of ER mutations. Our findings provide valuable information regarding PR action in the presence of E2-independent ER. Understanding the exact interplay between PR activity and ER function is crucial to optimizing treatment strategies for patients with advanced disease, particularly in the face of therapy resistance.

## RESULTS

### PR expression levels and protein interactions are altered in models of ESR1 mutations

For our studies, we utilized established T47D (24) and MCF7 (25) ER+ breast cancer models that express wild type (WT) ER or mutant ER (Y537S or D538G). Because PR is traditionally an E2-induced ER target gene, we sought to evaluate PR status in these breast cancer cell lines. Typically, ligand-activated ER induces robust PR expression. Accordingly, in T47D WT ER cells, total and phosphorylated PR (Ser294, Ser345) levels were increased following pre-treatment with E2 for 48h and subsequent treatment with synthetic progestin R5020 for 1h relative to cells expressing WT ER that did not receive E2 (**Fig. 1A**). As expected of constitutively active ERs, basal phosphorylated ER (Ser118) levels were increased in T47D Y537S and D538G ER+ cells relative to cells expressing WT ER. Notably, E2 was not needed to induce PR in T47D Y537S and D538G ER cells, which exhibited higher basal total and phosphorylated PR levels compared to cells expressing WT ER and upon R5020 treatment only (**Fig. 1A**). We also observed similar increases in basal PR levels within MCF7 models (**Supplementary Fig. 1**). These results show that total and phosphorylated PR levels are elevated in breast cancer cells expressing mutant ER in the absence of E2.

**Figure 1.**
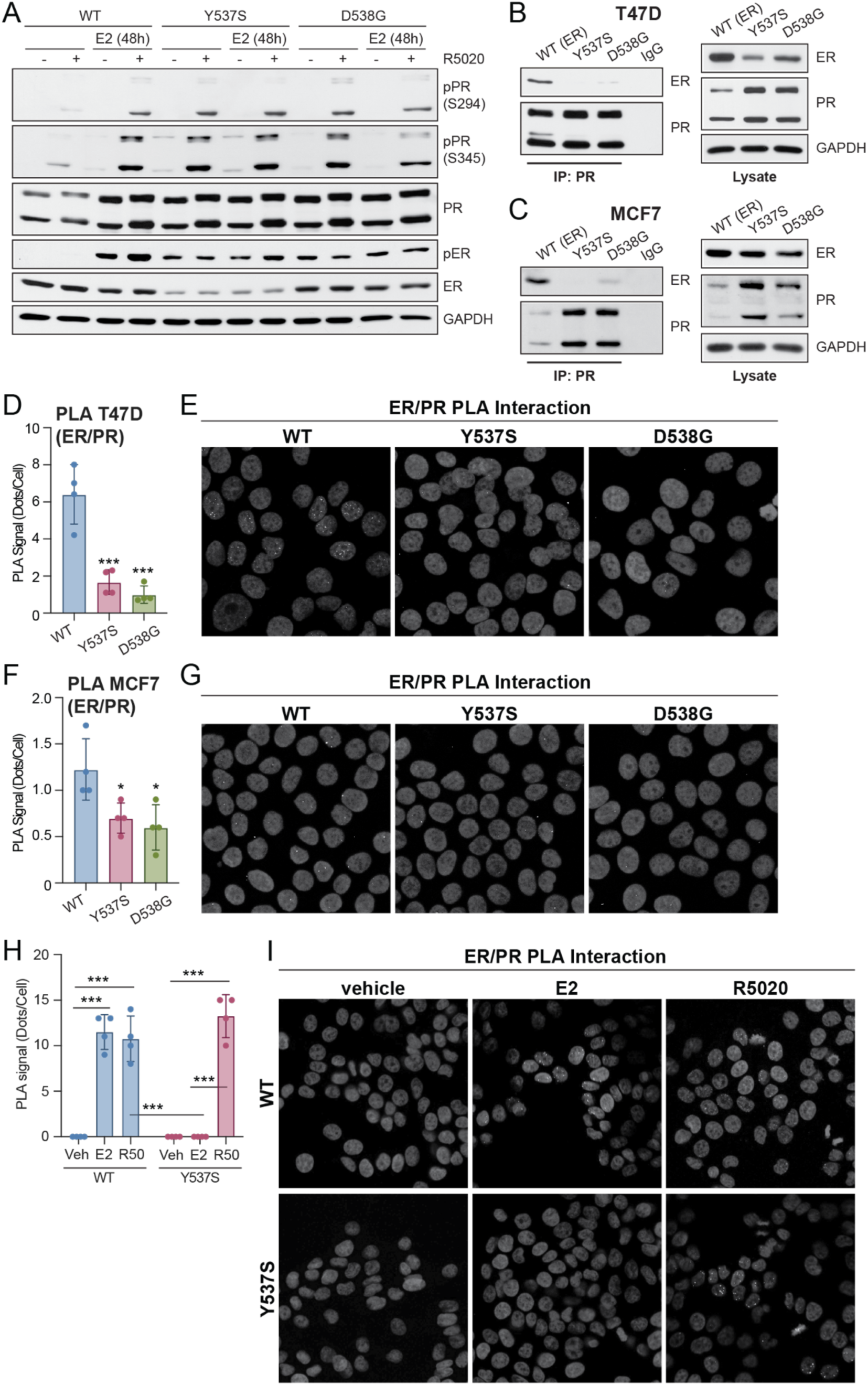
ER/PR interaction is altered in the context of ER mutations. (**A**) Western blot of T47D ER (WT, Y537S, D538G) cells pre-treated with E2 (1 nM) for 48 h followed by treatment with vehicle or R5020 (10 nM) for 60 min. Co-immunoprecipitation of ER and PR in (**B**) T47D and (**C**) MCF7 ER cells. Cell lysate controls (*right*). ER/PR interaction measured by proximity ligation assay (PLA) in (**D, E**) T47D and (**F, G**) MCF7 ER cells. (**H, I**) ER/PR interaction measured by PLA in T47D ER cells treated with vehicle, E2 (1 nM), or R5020 (10 nM) for 30 min. Quantification and representative images are shown for PLA. Graphed data represent the mean ± SD (n = 4), unless otherwise indicated. * p < 0.05, ** p < 0.01, *** p < 0.001.

PR has been shown to interact with ER through regulatory crosstalk (26) and direct mechanisms (12) to modify ER signaling and transcriptional programs. To evaluate if the ER/PR protein complex is altered in the context of ER mutations, we performed co-immunoprecipitation (co-IP assays). ER/PR interactions were present in T47D (**Fig. 1B**) and MCF7 (**Fig. 1C**) expressing WT ER but greatly reduced in both Y537S and D538G ER+ models in the absence of added hormone. Consistently, proximity ligation assays (PLA) in both T47D (**Fig. 1D, 1E**) and MCF7 (**Fig. 1F, 1G**) cells showed that basal ER/PR interactions were significantly reduced in cells expressing either Y537S or D538G ER. We next evaluated E2 or progestin (R5020)-induced ER/PR interactions. Cells expressing WT ER showed a significant increase in the ER/PR interaction following either E2 or R5020 treatment, whereas cells expressing mutant Y537S ER showed an increased ER/PR interaction when treated with progestin, but not E2 (**Fig. 1H, 1I**). Together, these results show that basal and hormone-regulated PR interactions with ER are altered in the context of *ESR1* mutations and that although mutant ERs are indeed E2-independent, progestin treatment still promotes ER/PR complexes.

### Breast cancer cells harboring ESR1 mutations have elevated stem-like cell populations

The expression of mutant ERs in breast tumors has been reported to promote CSC phenotypes (27). To build on these findings, we measured relative ratios of CD44^hi^/CD24^lo^ cells, a common marker of breast CSCs (28), in T47D and MCF7 cells cultured in 2D adherent or 3D tumorsphere conditions. Breast CSCs comprise a small portion of the overall tumor cell population, and 3D tumorsphere conditions enable us to expand CSCs *in vitro* (29). In 2D conditions (**Fig. 2A, Supplementary Fig. 2A, 2B**), Y537S ER exhibited the highest CD44^hi^/CD24^lo^ ratios (T47D, 16.2% ± 4.5; MCF7, 18.2% ± 0.7) compared to WT and D538G ER, and these trends were further enhanced in 3D culture conditions (T47D, 43.6% ± 2.0; MCF7, 38.7% ± 5.8) (**Fig. 2B, Supplementary Fig. 2C, 2D**). Notably, CD44^hi^/CD24^lo^ ratios were also significantly increased in D538G ER relative to WT in 3D conditions (**Fig. 2B**), a pattern that was not observed in 2D (**Fig. 2A**). Tumorsphere assays, which measure CSC activity and the property of CSC self-renewal, confirmed that T47D (**Fig. 2C, *top***) and MCF7 (**Fig. 2C, *bottom***) Y537S ER+ models had greater capacity for breast CSC formation. Additional E2 or R5020 hormone treatment did not increase tumorsphere forming ability in models expressing either WT or mutant ER (**Fig. 2D**). Treatment with endocrine therapies, fulvestrant (Fulv) (**Fig. 2E**) or tamoxifen (4OHT) (**Fig. 2F**), reduced tumorsphere formation in both WT and mutant ER+ models. We also tested next-generation endocrine therapies, elacestrant (30) and lasofoxifene (31,32), which have both been shown to be effective against mutant forms of ER. Elacestrant and lasofoxifene were effective in reducing tumorsphere formation in WT as well as mutant ER+ models (**Fig. 2F**). These results demonstrate that ER mutations confer enriched expansion of breast CSC populations, and that breast cancer cells expressing Y537S ER exhibit a stronger CSC phenotype overall relative to WT or D538G ER+ cells.

**Figure 2.**
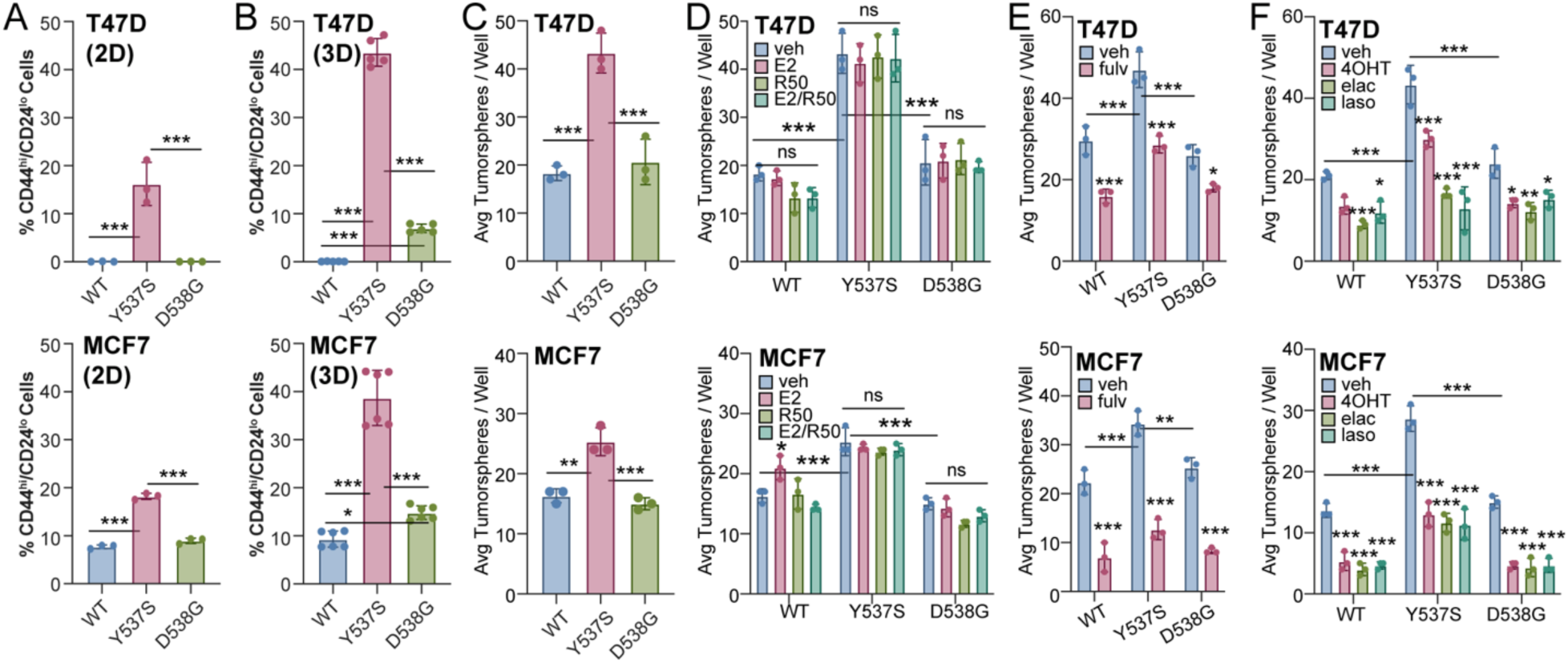
ER mutants display cancer stem cell phenotypes. CD44^hi^/CD24^lo^ populations in (**A**) T47D (*top*) and MCF7 (*bottom*) ER (WT, Y537S, D538G) cells cultured in 2D (adherent) conditions. CD44^hi^/CD24^lo^ populations in (**B**) T47D (*top*) and MCF7 (*bottom*) 3D (tumorsphere) conditions. Secondary tumorspheres in (**C**) T47D (*top*) and MCF7 (*bottom*) ER cells. Secondary tumorspheres in (**D**) T47D (*top*) and MCF7 (*bottom*) ER cells treated vehicle, E2 (1 nM), and/or R5020 (10 nM). Secondary tumorspheres in (**E, F**) T47D (*top*) and MCF7 (*bottom*) ER cells treated with vehicle, fulvestrant (fulv; 100 nM), 4-hydroxytamoxifen (4OHT; 100 nM), elacestrant (elac; 100 nM), and lasofoxifene (laso; 100 nM). Graphed data represent the mean ± SD (n = 3), unless otherwise indicated. * p < 0.05, ** p < 0.01, *** p < 0.001.

We next evaluated the impact of modulating PR on biological phenotypes in models of *ESR1* mutations. Treatment with well-characterized antiprogestins (onapristone, RU486, or PRA-027) did not have a significant effect on tumorsphere formation in our T47D models (**Fig. 3A**), and only a modest effect was observed in MCF7 cells expressing WT ER or the Y537S or D538G *ESR1* mutation (**Fig. 3B**). Because we did not observe robust reduction in tumorsphere formation by antiprogestins, we experimentally manipulated PR levels by genetic knockdown using shRNA in T47D WT and Y537S ER+ cells to assess the effect of PR knockdown on advanced cancer cell phenotypes. Since cells expressing Y537S ER exhibited the most significant effects in CSC assays compared to cells expressing D538G ER (**Fig. 2**), we proceeded with Y537S ER. Western blots showed that PR expression levels were greatly reduced following lentiviral expression of shRNA targeting PR in both T47D WT and Y537S ER+ cells (**Fig. 3C**). Control studies with E2 treatment for 24 or 48 hours in shPR cells confirmed PR levels were not induced by E2 and that PR knockdown was successful (**Supplementary Fig. 3**). Tumorsphere assays in control (shGFP) and shPR cells demonstrated that PR knockdown significantly reduced tumorsphere formation in T47D cells expressing WT ER by 56.5% (shGFP: 15.3 ± 2.5; shPR: 6.7 ± 0.6) and Y537S ER by 77.1% (shGFP: 55.3 ± 1.5; shPR: 12.7 ± 1.5) (**Fig. 3D**). We expanded these studies to soft agar colony formation assays which measure anchorage independent growth. T47D WT ER+ cells have increased colony formation upon E2 treatment, whereas Y537S ER+ cells have significantly increased overall colonies compared to WT ER but did not respond to E2 (**Fig. 3E**). In both WT and Y537S ER+ models, PR knockdown also significantly reduced colony formation as measured by soft agar anchorage independent growth assays (**Fig. 3E**).

**Figure 3.**
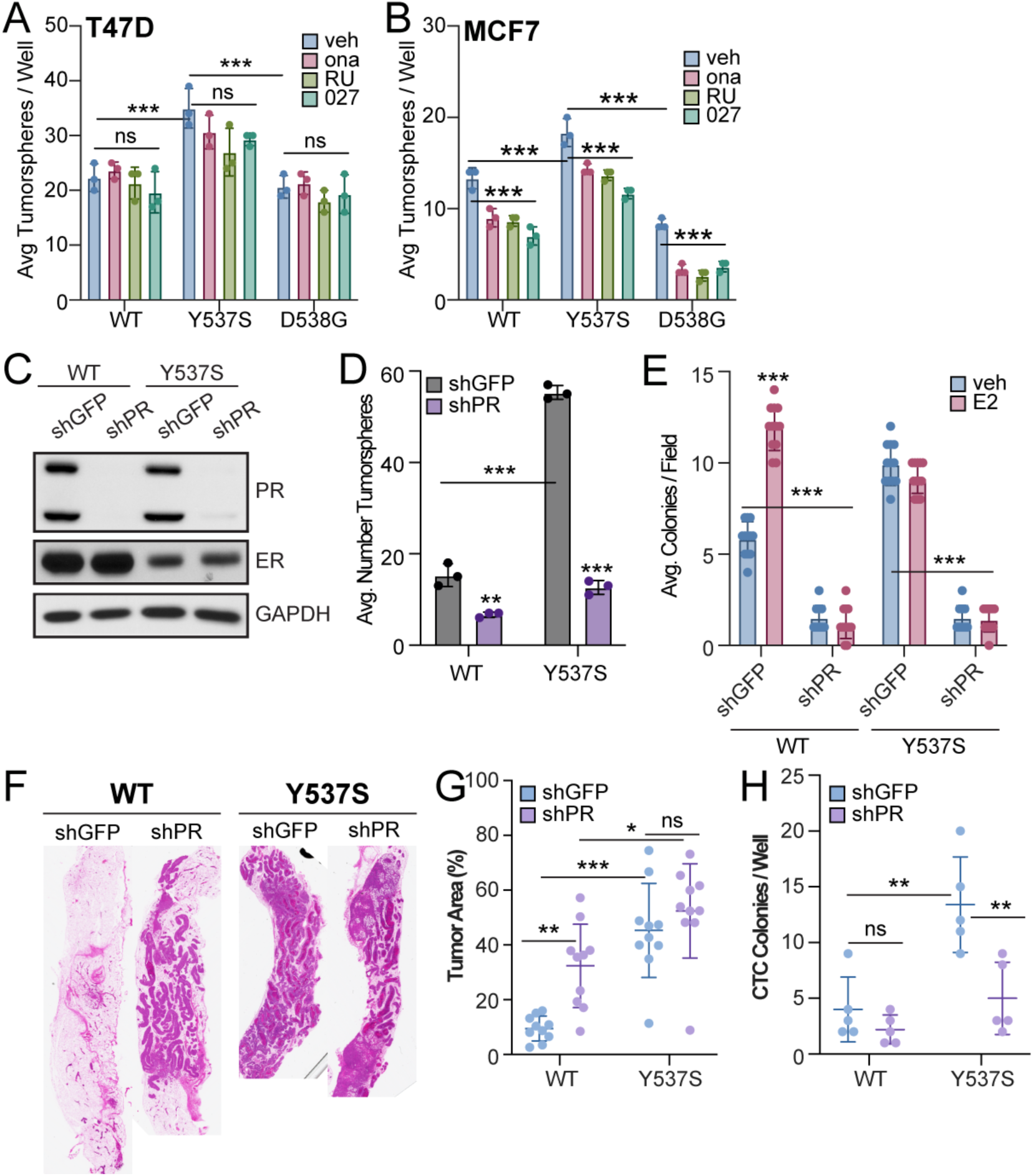
PR knockdown affects cancer cell biology phenotypes in ER models. Secondary tumorspheres in (**A**) T47D and (**B**) MCF7 ER cells treated with veh (EtOH), onapristone (200 nM), RU486 (200 nM), or PRA-027 (200 nM). (**C**) Western blot of T47D ER cells with PR knockdown. (**D**) Secondary tumorspheres in T47D ER PR knockdown cells. (**E**) Soft agar colony formation in T47D ER PR knockdown cells treated with veh (EtOH) or E2 (1 nM). (**F**) Representative H&E stains from T47D WT and Y537S ER (shGFP, shPR) MIND glands. (**G**) Tumor area (%) calculated from H&E sections from T47D WT and Y537S ER (shGFP, shPR) MIND glands. (**H**) Average number of colonies/well (CTCs). Graphed data represent the mean ± SD (n = 3), unless otherwise indicated. * p < 0.05, ** p < 0.01, *** p < 0.001.

To examine the *in vivo* relevance and impact of PR in the context of *ESR1* mutations, we utilized the mouse mammary intraductal (MIND) xenograft model (33,34) whereby tumor cells are directly injected into the mammary gland ducts. This approach preserves the luminal phenotype of the cells, unlike the traditional mammary fat pad approach in which injected ER+ breast cancer cells require supplemental estrogen to form tumors and acquire basal characteristics (34). T47D WT and Y537S ER+ cells harboring control (shGFP) or PR knockdown (shPR) were injected into the inguinal mammary glands of adult female immunocompromised mice. Mice were sacrificed at 8-weeks post injection and tumor area (%) was calculated from hematoxylin and eosin (H&E) images (**Fig. 3F**). These analyses revealed that the mean tumor volume differences between mice injected with cells expressing WT ER shGFP (9.5% ± 4.5) and Y537S ER shGFP (45.3% ± 17.2) was significant (*p* < 0.0001, **Fig. 3G**). Surprisingly, knockdown of PR in T47D WT ER cells significantly increased primary tumor burden (shPR = 32.3% ± 15.2, *p* = 0.006) while we observed no detectable effects in Y537S ER+ cells (shPR = 52.4% ± 17.2, *p* = 0.694) compared to their respective shGFP controls (**Fig. 3G**). To assess disseminated tumor cells, blood was also collected by exsanguination at the time of euthanasia, processed to eliminate red blood cells and then seeded into soft agar assays to evaluate the presence of circulating tumor cells (CTCs) (15). We observed a significantly higher number of CTC colonies from the peripheral blood of mice injected with Y537S ER shGFP (13.4 ± 4.3) relative to that observed in WT ER shGFP controls (4.0 ± 2.9) (**Fig. 3H**). Notably, PR knockdown significantly reduced the number of CTCs in mice injected with Y537S ER shPR cells (5.0 ± 3.2, *p* = 0.003) but not in mice injected with WT ER shPR cells (2.2 ± 1.3, *p* = 0.799) relative to the respective shGFP controls (**Fig. 3H**). These xenograft results suggest that PR expression increases cancer stem-like characteristics and metastatic potential as measured by disseminated CTCs in the context of *ESR1* mutations.

### PR regulates distinct transcriptomes and cistromes in ESR1 mutant breast cancer models

We next sought to examine the transcriptional consequence of PR activity in the context of *ESR1* mutations. Our studies demonstrate response to progestin regarding ER/PR complex formation, but not estrogen in cells harboring Y537S ER (**Fig. 1H**). We performed RNA-seq studies in T47D ER+ models grown in 2D (adherent) or 3D (tumorsphere) conditions. For 2D studies, T47D WT and Y537S ER+ cells were treated with vehicle (ethanol) or E2 for 48 hours (i.e., to induce PR expression), followed by R5020 treatment for 6 hours. PR knockdown (shPR) models were included as controls. Hierarchical clustering (**Supplementary Fig. 4A**) and principal component analysis (PCA) (**Fig. 4A**) demonstrated a robust response to R5020 in treated samples for both WT and Y537S ER+ models. Gene expression changes in response to hormone treatment (E2 and/or R5020) were quantified and plotted as cross plots for WT (**Fig. 4B**) and Y537S ER (**Fig. 4C**). Notably, R5020 treatment was a major driver of transcriptional response between cells harboring either WT ER or Y537S ER, represented by the differentially expressed genes (DEGs) shown as blue dots. E2 treatment drove changes in transcription for WT ER (i.e., pink dots), with some shared DEGs regulated in the same direction by either E2 or R5020 treatment denoted as yellow dots. Conversely, Y537S ER showed few transcriptional differences with E2 treatment (i.e., pink dots) and no E2 or R5020 shared DEGs (i.e., yellow dots) (**Fig. 4C**). Consistent with these cross plots, Venn diagram analysis of DEGs revealed similar findings. In sum, both WT and Y537S ER+ cells were responsive to R5020 treatment in the absence of E2 (**Fig. 4D**), whereas only cells expressing WT ER responded to E2 alone (**Supplementary Fig. 4B**). Cells expressing WT or Y537S ER also exhibited differential responses to combined treatment with both E2 and R5020 (**Supplementary Fig. 4C**). We next performed comparative gene set enrichment analysis (GSEA) using GO ontology terms on DEGs in WT and Y537S ER+ models and in response to R5020 treatment (**Fig. 4E**). This analysis revealed that the R5020-induced response modulated distinct biological pathways and cellular functions in breast cancer cells expressing WT ER compared to cells harboring Y537S ER. Specifically, R5020-induced WT ER-enriched pathways included ERK1/2 cascades and those associated with basal plasma membrane function linked to proliferation. Conversely, R5020-induced Y537S ER-enriched pathways correlated to migration and metastasis such as actin fiber structure and assembly. Together, these analyses support a role for PR activation in more aggressive cancer phenotypes associated with Y537S *ESR1* mutations.

**Figure 4.**
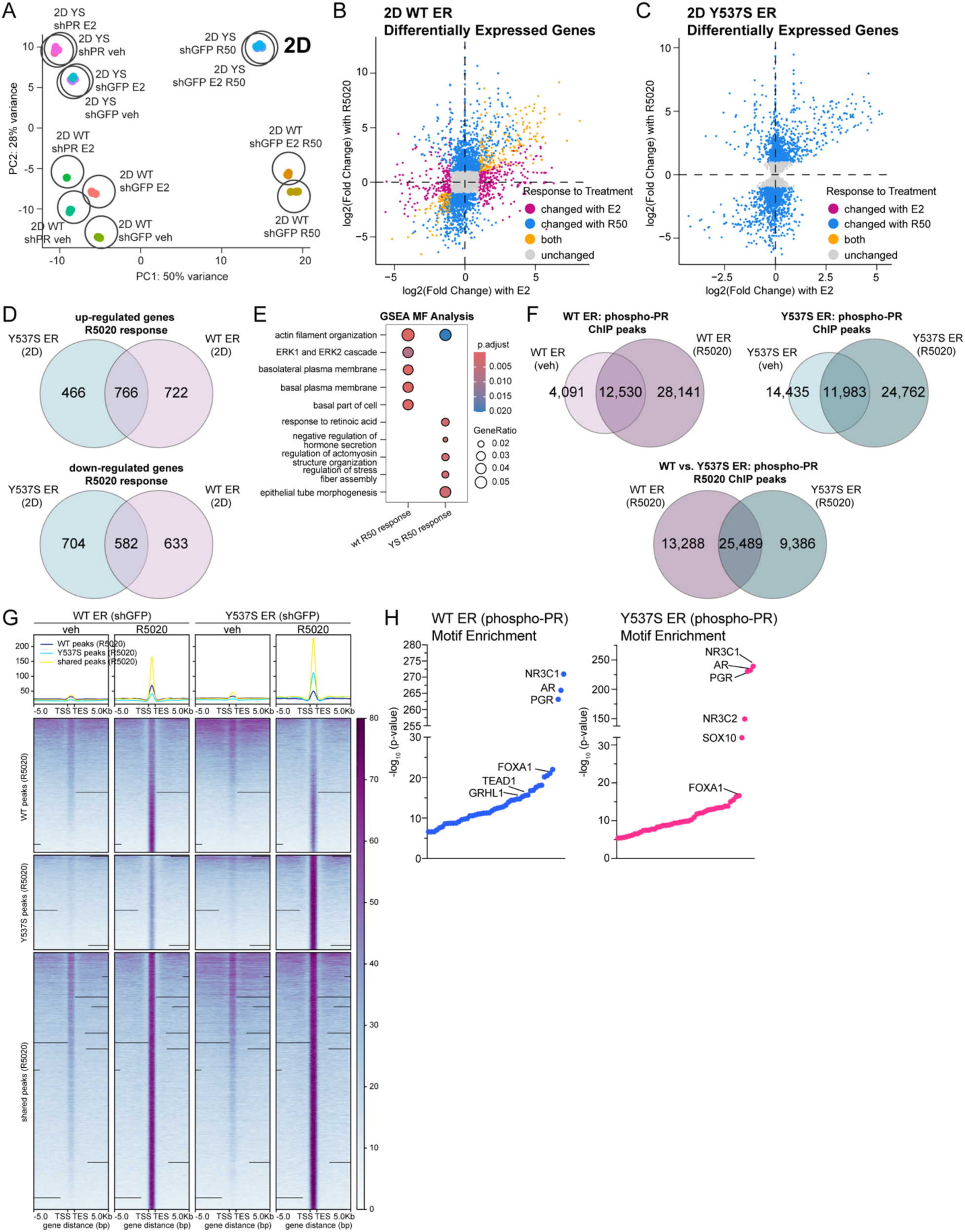
WT and Y537S ER have distinct transcriptomes and genomic binding patterns in response to progestin in 2D culture conditions. For RNA-seq studies, T47D WT and Y537S (shGFP) cells were treated with vehicle, E2 (1 nM), R5020 (10 nM), or combination for 6h. T47D WT and Y537S (shPR) cells are included as controls. (**A**) PCA plot showing distribution of 2D sample groups treated with hormones. Cross plots showing differentially expressed genes for (**B**) WT and (**C**) Y537S shGFP in response to E2, R5020, and combination treatment. (**D**) Venn diagrams showing differentially expressed genes up or downregulated >2-fold in response to R5020. (**E**) GSEA analysis dot plot showing enriched pathways in WT and Y537S ER in response to R5020. For ChIP-seq studies, T47D WT and Y537S (shGFP) cells were treated with vehicle or R5020 (10 nM) for 1h. (**F**) Venn diagrams represent peak overlap analysis for phospho-PR in response to R5020 in WT ER and Y537S ER, respectively (*top*). *Bottom:* Venn diagram shows overlap analysis for phospho-PR in R5020 treatment condition for WT versus Y537S ER. (**G**) Density plots show distinct and shared peaks in WT and Y537S ER in response to R5020. (**H**) Homer motif analysis for phospho-PR in WT ER (*left*) and Y537S ER (*right*) to highlight top transcription factor motifs enriched near phospho-PR binding sites.

We next performed ChIP-seq studies using a custom phospho-Ser294 PR (pSer294) antibody to determine if ER mutation influences PR genomic occupancy. PR Ser294 is an N-terminal residue within a proline-directed consensus phosphorylation site for MAPK or CDKs, and previous studies from our lab have shown that it is required for endocrine (i.e., tamoxifen) resistance and CSC phenotypes (18,20). Phospho-Ser294 PR exhibits altered target gene selection at a subset of PR-and ER-regulated genes (20). Peak overlap analysis revealed that phospho-PR exhibited some PR binding at distinct sites in the vehicle-treated samples but significantly increased in response to R5020 treatment in both WT and Y537S ER+ models as shown by their respective Venn diagrams (**Fig. 4F, *top***). Comparison of R5020-induced phospho-PR cistromes revealed 25,489 shared peaks between cells expressing WT or Y537S ERs; however, phospho-PR had 13,288 unique binding sites whereas it gained 9,386 new sites in the Y537S ER background (**Fig. 4F, *bottom***). Density plots illustrate the R5020-dependent increase in ChIP-seq signal and distinct pSer294-PR binding sites between WT and Y537S ER (**Fig. 4G**) and respective controls (input and shPR) (**Supplementary Fig. 5**). Motif analysis revealed transcription factor motifs enriched at pSer294-PR binding sites in cells expressing WT ER compared to Y537S ER (**Fig. 4H**), suggesting that altered co-factor recruitment may in part contribute to the changes in transcriptional gene programs observed from our RNA-seq studies. Taken together, these data illustrate that gene regulation and genomic binding elicited by progestin-induced responses is distinct in breast cancer cells harboring *ESR1* mutations relative cells expressing WT ER.

### PR-driven CSCs exhibit changes in key transcriptional programs in ESR1 mutant models

Our RNA-seq studies from cells grown in 2D culture showed that activated PR induced detectable changes in transcription and genomic binding (**Fig. 4**). Although our studies showed that antiprogestins had modest effects (**Fig. 3A, 3B**), we observed a striking reduction in tumorspheres when PR was knocked down (WT ER, 56.5% decrease; Y537S ER, 77.1% decrease) (**Fig. 3D**). Therefore, we chose to perform RNA-seq on T47D ER models with or without PR knockdown (shGFP or shPR) grown in 3D tumorsphere conditions to identify differences unique to CSC-enriched samples. Hierarchical clustering of DEGs demonstrated that Y537S ER induced a distinct transcriptional response compared to WT ER (**Fig. 5A**), which was consistent with PCA plots where Y537S ER shGFP was the largest driver of variance between samples (80%) (**Fig. 5B**). We next compared the log2 fold change in DEGs relative to WT shGFP across conditions. This analysis further illustrated the dependence on the presence of PR for Y537S ER-induced target genes compared to WT ER and revealed that many Y537S-induced genes return to baseline expression levels upon PR knockdown (**Fig 5C**). Venn diagram analysis illustrated DEGs up or downregulated >2-fold by PR knockdown (shPR versus shGFP comparison; Y537S shGFP = 2107 up-regulated, 862 down-regulated and WT shGFP = 270 up-regulated, 589 down-regulated) (**Fig. 5D**). There are fewer overlapping genes between Y537S and WT ER in this comparison (90 total; 60 up-regulated, 30 down-regulated). We next performed comparative GSEA using GO ontology terms to evaluate pathways impacted in 3D culture conditions. Specifically, WT ER shGFP exhibited suppression of pathways involved in the immune response (**Fig. 5E**), whereas Y537S ER shGFP exhibited activation of unfolded protein binding (**Fig. 5F**) when compared across all other conditions. Follow-up analysis of the DEGs from Y537S ER samples (shPR versus shGFP comparison) using the Ingenuity Pathway Analysis (IPA) canonical pathways predicted increases in the unfolded protein response (UPR) and other related stress-induced pathways to UPR (e.g., autophagy, cholesterol biosynthesis, and endoplasmic reticulum stress) (**Fig. 5G**). Lastly, we performed GSEA analysis using the MSigDB Hallmarks gene set to further emphasize negative enrichment of immune response pathways (e.g., interferon gamma response, alpha gamma response) in WT ER (**Fig. 5H, *left***), and positive enrichment of the UPR and related cellular stress-induced signaling pathways (e.g., hypoxia) in Y537S ER samples (**Fig. 5H, *right***). Together, these results show that the PR-driven pathways identified from our 3D RNA-seq studies are clearly different between WT and Y537S ER. Moreover, our findings also indicate that CSC-enriched populations (i.e., enabled by 3D culture) elicit a distinct transcriptional response when compared to our 2D RNA-seq studies.

**Figure 5.**
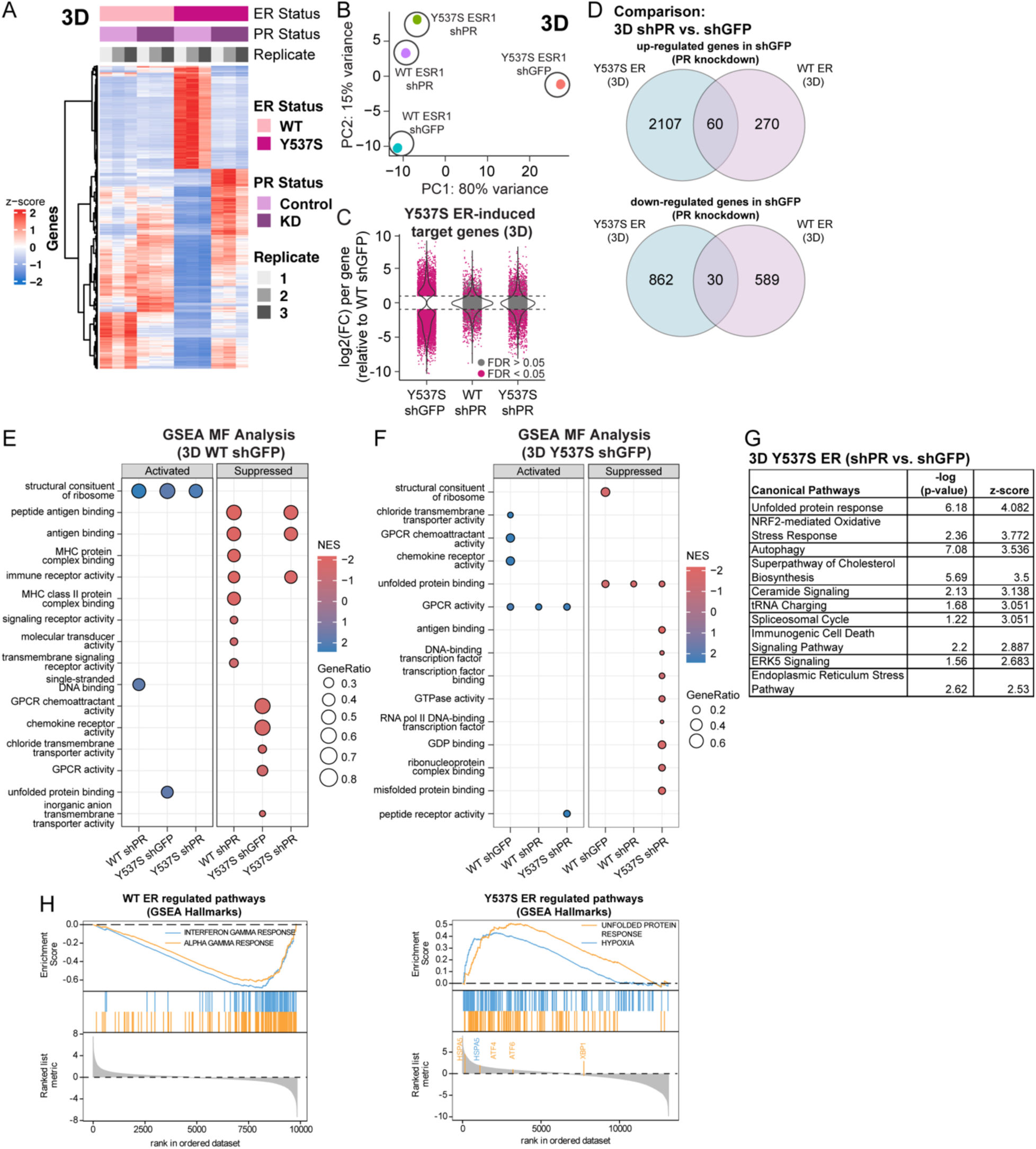
PR knockdown in Y537S ER remodels the transcriptome in 3D culture conditions. T47D WT and Y537S (shGFP, shPR) cells were cultured in 3D tumorsphere conditions. (**A**) Hierarchal clustered heat map of differentially expressed genes (DEGs) in 3D samples. (**B**) PCA plot showing distribution of 3D sample groups. (**C**) Comparison of log2 (fold change) of Y537S ER-induced target genes across conditions in 3D tumorsphere samples. Many Y537S-induced DEGs are no longer altered upon PR knockdown. (**D**) Venn diagrams showing differentially expressed genes up or downregulated >2-fold in PR knockdown cells (3D shPR versus 3D shGFP). GSEA analysis dot plot showing activated and suppressed pathways upon PR knockdown in (**E**) WT and (**F**) Y537S ER. (**G**) Ingenuity Pathway Analysis (IPA) showing canonical pathways in 3D T47D Y537S ER. (**H**) GSEA enrichment plots showing selected WT ER (*left*) and Y537S ER (*right*) regulated pathways enriched upon PR knockdown.

### The UPR is a targetable PR-driven CSC pathway in the context of ESR1 Y537S mutation

The UPR is a cellular stress response that regulates protein homeostasis and protects cells when misfolded or unfolded proteins build up in the endoplasmic reticulum. This pathway is also highly correlated with endocrine resistance (35). Studies using ER+ breast cancer cell lines have demonstrated that ER mutations (Y537S, D538G) upregulate the UPR independently of E2 and elevate selected markers of UPR activation upon progestin treatment (24); these studies were performed in 2D culture conditions. Our 3D RNA-seq studies suggest that the UPR pathway is also modulated by PR in cells harboring Y537S ER and cultured in CSC-enriched conditions (**Fig. 5**). Activation of the UPR occurs through separate signaling arms that are controlled by different sensors (IRE1, ATF6, and PERK). To assess which signaling arm impacts PR-dependent CSC populations in ER mutations, we performed tumorsphere assays with T47D and MCF7 models treated with inhibitors specific to each UPR arm (KIRA8, inhibitor of IRE1; Ceapin-A7, inhibitor of ATF6; and GSK2606414, inhibitor of PERK). All UPR inhibitors tested exhibited a relatively modest effect in both WT and Y537S ER+ models, with the highest levels of tumorsphere reduction reaching 40.7% (GSK2606414) in T47D (**Fig. 6A**) and 45.0% (KIRA8) in MCF7 (**Fig. 6B**) models when high micromolar concentrations of inhibitors were used. Additionally, treatment with GSK2606414, but not KIRA8 or Ceapin-A7, showed a modest diminution of R5020-induced phospho-PR levels in Y537S ER+ cells (**Supplementary Fig. 6A-6C**). Mild UPR activation in ER+ breast cancer can be protective and acts as a mechanism for cells to handle increased protein folding capacity; conversely, hyperactivation of the UPR reverses this protection (36). To leverage this, we used ErSO, a small molecule UPR hyperactivator that has been shown to reduce tumor progression and metastasis in ER+ mouse xenografts by converting the UPR from a cytoprotective pathway to one that is cytotoxic (37). Treatment with ErSO was highly effective at reducing tumorsphere formation (**Fig. 6C**) and blocked R5020-driven PR Ser294 phosphorylation in T47D WT and Y537S ER+ models (**Fig. 6D**). These studies indicate that blocking specific UPR signaling arms may not be an effective way to reduce PR-driven CSC phenotypes that are enhanced in Y537S *ESR1* mutations, whereas hyperactivation of the UPR pathway with compounds such as ErSO impacts both CSC biology and PR activity.

**Figure 6.**
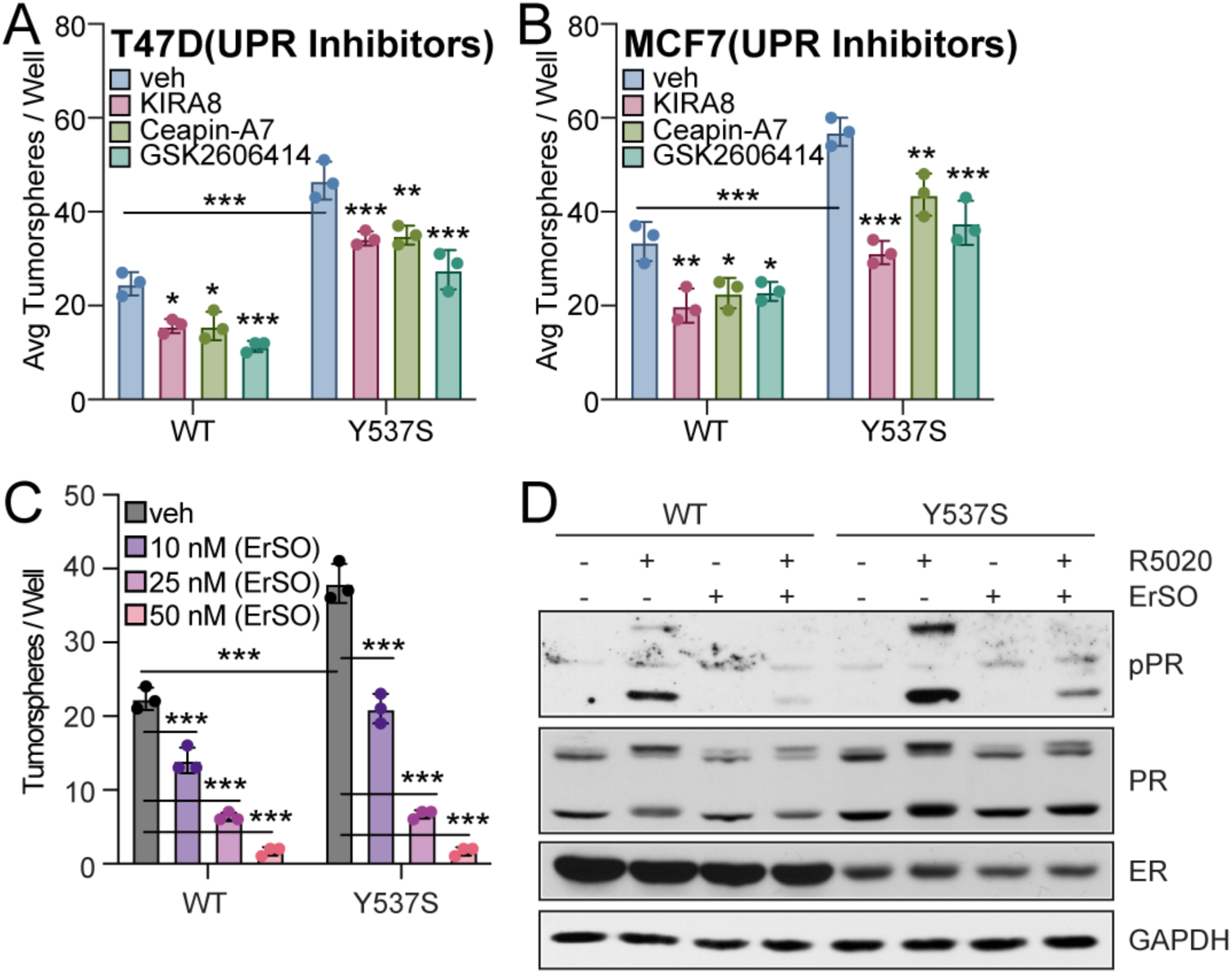
UPR is a potential PR-mediated CSC pathway. Primary tumorspheres in (**A**) T47D and (**B**) MCF7 ER cells treated with vehicle, KIRA8 (5 µM), Ceapin-A7 (5 µM), or GSK2606414 (5 µM). (**C**) Secondary tumorspheres in T47D ER cells (WT, Y537S) treated with ErSO (10, 25, or 50 nM). (**D**) Western blot of T47D ER cells (WT, Y537S) pre-treated with ErSO (50 nM; UPR activator) for 1h followed by vehicle (EtOH) or R5020 (10 nM) for 1h. Graphed data represents mean ± SD (n = 3). * p < 0.05, ** p < 0.01, *** p < 0.001.

## DISCUSSION

In this study, we demonstrate that PR action is altered in the context of *ESR1* gene mutations that create constitutively active (i.e., Y537S) ERs. Response to progesterone treatment is distinct in breast cancer cells harboring Y537S ER, despite its reduced E2 sensitivity relative to WT ER. Our findings show that PR activity is required for enhanced breast CSC phenotypes observed in Y537S ER, and 3D RNA-seq studies identified PR-dependent pathways in the context of activated but E2-independent ER that can be pursued to specifically target CSC populations in endocrine resistant settings.

Improved treatment options are urgently needed for metastatic ER+ breast cancer patients whose cancers have progressed following standard endocrine therapies. Major efforts have focused on developing next-generation SERMs and SERDs (6) capable of targeting ER in breast cancer cells harboring *ESR1* mutations. To date, elacestrant and imlunestrant are the only FDA approved SERDs for advanced metastatic breast cancer patients that show activity against *ESR1* mutations. Other SERMs and SERDs have shown clinical promise in *ESR1* mutant settings, such as giredestrant (38) and lasofoxifene (39). Drug development efforts are primarily focused on ER-targeted approaches, and it will be crucial to consider other central drivers such as PR. Increasing evidence supports PR as having a key role in ER+ breast cancer progression, especially in disseminated CTC and/or CSC populations. For example, ER/PR crosstalk is one mechanism known to modulate ER signaling and transcriptional events in 2D culture conditions (26). A recent study from the Greene laboratory reported that the ER/PR interaction is altered in the presence of Y537S ER, and identified four genes that ER and PR have shared regulatory binding sites for, including insulin receptor substrate-1 (IRS-1) (40). Studies from the Lange lab also report a link between PR and IRS-1, and show that a phosphorylated PR (specifically, the PR-B isoform) is needed to cooperate with IRS-1 to promote outgrowth of tamoxifen-resistant and stem-like breast cancer cells (18). Phospho-Ser294 PR is found in ∼50% of ER+ breast cancers (20) and has more recently been associated with fallopian tube STIC lesions known to give rise to high grade serous ovarian cancers (41). In breast cancer models, phospho-Ser294 PR target genes correlate with poor outcome in breast cancer patients (i.e., public data), and these phosphorylated PR species have been shown to be enriched in ER+ PDX metastases in the bone and/or brain of implanted NSG mice (18). Thus, routine targeting of activated PRs (i.e., phosphorylated species) as part of ER-targeted therapies may be a necessary approach to overcome recurrence of ER+ breast cancers.

The results of this study show that the Y537S ER mutation significantly alters protein-protein interactions with PR (**Fig. 1**) and transcriptional responses (**Fig. 4**) following hormone treatments. WT ER responds to E2 and progestin alone or in combination. Conversely, Y537S ER only exhibits a response to progesterone treatment, and this is largely distinct from the transcriptional gene programs identified in cells expressing WT ER (**Fig. 4B, 4C**). ChIP-seq studies performed with pSer294 PR antibodies provided additional insight into how WT and Y537S ER expression influence the genomic occupancy of PR. In the absence of R5020 treatment, more basal pSer294-PR peaks were detected in Y537S ER cells (14,435 unique peaks, veh) compared to WT ER cells (4,091 unique peaks, veh) (**Fig. 4F**). This suggests that there is increased ligand-independent genomic activity of phospho-PRs in the context of Y537S ER, which may account for some differences observed in basal transcriptional activity between these cells (**Supplementary Fig. 4B**). R5020 treatment induced p-PR binding in both WT ER cells (28,141 unique peaks, R5020) and Y537S ER+ cells (24,762 unique peaks, R5020) with a large degree or overlap in these cistromes (**Fig. 4F**). A separate study reported that ER mutations enhance progesterone response through cooperation with PR; however, these ChIP-seq were performed looking using total PR antibodies (42). Our findings suggest that it will be important to distinguish between ligand-dependent (i.e., R5020) and ligand-independent activation of PR to appropriately target these phosphorylated PR species in mutant ER tumors. In this study, we evaluated contributions from both PR isoforms, because current antibodies used in routine clinical testing do not distinguish between isoforms. The levels of PR-A and PR-B isoforms are typically balanced (1:1 ratio) in normal mammary epithelial cells; this ratio becomes unbalanced in breast tumors (43). Studies from our lab and others have shown that PR isoforms differentially regulate transcriptional outcomes (15,44) and exhibit unique genomic bindings patterns (15). Future studies to dissect isoform-specific PR functions in the context of ER mutations and in 3D contexts will further inform efforts to therapeutically target total and phosphorylated PR.

PRs are emerging as major drivers of CSC biology in ER+ breast cancer, in both therapy-responsive and -resistant settings. Studies have shown increased CSC activity following treatment with endocrine therapy (e.g., letrozole, tamoxifen, or fulvestrant) in breast cancer cell models and tumor samples (45,46). Our lab has shown that PR knockdown or co-targeting with antiprogestins effectively reduces tumorsphere formation in tamoxifen-resistant models (18). In models of Y537S *ESR1* mutation, we demonstrate that PR knockdown has a significant effect on reducing CSC phenotypes (**Fig. 3D**) and metastatic potential (**Fig. 3H**). Cells expressing Y537S ER had considerably more genes impacted by PR knockdown (2969 total genes) compared to cells expressing WT ER (859 total genes) in 3D culture conditions (**Fig. 5D**). Notably, the UPR was identified as a candidate pathway in response to PR knockdown in Y537S ER+ models in our 3D RNA-seq studies (**Fig. 5**). Other studies have shown enhanced UPR activation in breast cancer cells harboring Y537S ER; these studies were performed in 2D culture conditions (24). Our 3D studies further distinguish that this response is enhanced in CSC-enriched samples and is a PR-dependent pathway. We show UPR inhibitors that target specific arms of the pathway exhibited modest effects in reducing CSC populations and demonstrated that hyperactivation of the UPR pathway with ErSO is a much more effective approach (**Fig. 6**). It is important to clarify that our studies primarily focused on Y537S ER given its relatively common occurrence in women taking long-term aromatase inhibitors (6) and more pronounced CSC phenotype; however, follow-up studies focusing on PR-driven CSC biology in D538G ER+ models are warranted to evaluate commonalities and differences between each respective ER mutation and its functional consequences with regard to PR or other related steroid hormone receptors. Additionally, further dissection of phosphorylated PR species (Ser294 versus Ser345 or Ser81 sites) is needed in cells harboring ER mutations given its significant role in CSC biology. This information would help guide how to target phospho-PR-driven actions in different states of endocrine resistance. The approaches taken to target non-CSCs compared to CTCs/CSCs will undoubtedly need to differ, and a better understanding the alterations that occur specifically in breast CSCs is needed to effectively reduce expansion of this resistant cell population. Recent studies performed in a cohort of 24 premenopausal women with increased breast cancer risk demonstrated that the antiprogestin ulipristal acetate induced mammary stromal remodeling coupled to luminal progenitor suppression (47). Although we also evaluated antiprogestins in our studies (**Fig. 3A, 3B**), they showed only a modest effect in reducing breast CSC phenotypes in the context of ER mutations, likely because phosphorylated PR species tend to be resistant to antagonists with partial-agonist activity (20). Development of selective antiprogestins for clinical use has been limited (48), as evidenced by the fact that few selective PR ligands exist and no selective PR-targeted therapies (e.g., selective progesterone receptor modulators [SPRMs] or PROTACs) have been approved to date for treatment of ER+ breast cancers (49). Additionally, although clinical trials in other cancers have been carried out, there have been no trials to date that have assessed the therapeutic benefit of selectively targeting CSCs in ER+ breast cancer patients (50). As more evidence that PRs are important drivers of CSC expansion comes to light, these limitations underscore the importance of our work and why efforts to develop phosphorylated PR and CSC-specific targeting approaches are urgently needed to overcome resistant ER+ tumors, including *ESR1* mutant breast cancers.

## ACKNOWLEDGEMENTS

This work was supported by resources and staff at the University of Minnesota Genomics Core (UMGC), University Imaging Centers (UIC), and the University Flow Cytometry Resource (UFCR). We would like to acknowledge Fatemah Iman Dewji (University of Minnesota) for technical assistance with histological data analysis, Angela Spartz (University of Minnesota) for critical reading of this manuscript, David J. Shapiro (University of Illinois Urbana-Champaign) for gifting T47D *ESR1* cells (WT, Y537S, D538G) and ErSO compound, and Ben Ho Park (Vanderbilt University) for gifting MCF7 *ESR1* cells (WT, Y537S, D538G).

## MATERIALS AND METHODS

### General Reagents

Estradiol (E2; Sigma), R5020 (Sigma), onapristone (ona; Arno Therapeutics), 4-hydroxytamoxifen (4OHT; Sigma), RU486 (Sigma), hydrocortisone (Sigma), and fulvestrant/ICI 182,780 (ICI; Tocris) stocks were prepared in ethanol (EtOH). ErSO (David Shapiro, University of Illinois Urbana-Champaign), KIRA8 (Selleckchem), Ceapin-A7 (Selleckchem), GSK2606414 (Selleckchem), PRA-027 (Geoffrey Greene, University of Chicago), elacestrant (elac; MedChemExpress), lasofoxifene (laso; Sigma) stocks were prepared in DMSO. Epidermal growth factor (EGF; Sigma) was prepared in 0.1% BSA.

### Cell Culture

STR authentication for cell lines was performed by ATCC (August 2021). All cell lines were tested for mycoplasma prior to the initiation of experiments. T47D ESR1 cells (24) were cultured in phenol-red free MEM (Gibco, #51200038) containing either 10% FBS (for WT ER) (Corning) or 10% DCC (Y537S ER, D538G ER) (dextran coated charcoal-stripped FBS; Corning), 1% penicillin-streptomycin (Gibco), and 1% GlutaMax (Gibco). MCF7 ESR1 cells (25) were cultured in DMEM (Corning, #10013CV) containing 5% FBS and 1% penicillin-streptomycin. For experiments with hormone treatment (i.e., E2 or R5020), cells were hormone-starved in phenol-free modified IMEM (Gibco, # A1048801) containing 5% DCC for 16 hours prior to treatment. For 3D (tumorsphere) conditions, cells were grown in a phenol-red free DMEM/F12 (Corning, #16405CV) containing 0.5% methylcellulose (Sigma), 1% B27 supplement (Gibco), 1% penicillin-streptomycin, 5 μg/ml insulin (Gibco), 20 ng/ml EGF, 1 ng/ml hydrocortisone, 100 μM β-mercaptoethanol (MP Biomedicals) in ultra-low attachment (ULA) plates (Corning).

### Stable Cell Line Generation

Stable shPR (Sigma; clones TRCN0000003321, −3324) cells were created by transducing T47D cell lines with pLKO.1 lentivirus and maintained as described above with 0.5 µg/ml puromycin (MP Biomedicals).

### Cell Lysate Preparation

Cells were harvested in RIPA-lite lysis buffer [150 mM NaCl, 6 mM Na_2_HPO_4_, 4 mM NaH_2_PO_4_, 1 mM EDTA, 1 mM NaF, 1% Triton-X 100, 1X complete mini protease inhibitors (Roche), 1X PhosSTOP (Roche), and supplemented with 1 mM PMSF, 1 mM NaF, 0.5 mM Na_3_VO_4_, 10 mM beta glycerophosphate (BGP), and 20 µg/ml aprotinin].

### Immunoblotting

Antibodies used: phospho-PR (Ser294 [custom], Ser345 [custom]; Pierce), PR (D8Q2J, Cell Signaling), phospho-ERα (Ser118 [16J4], Cell Signaling), ERα (F-10, Santa Cruz Biotechnology), GAPDH (0411, Santa Cruz Biotechnology), goat anti-rabbit IgG-HRP (BioRad), and goat anti-mouse IgG-HRP (BioRad). Blots were developed with ECL using Super Signal West Pico Plus Chemiluminescence Substrate (Pierce) and imaged by film.

### CD44/CD24 Populations (Flow Cytometry)

For CD44/CD24 detection, cells were dissociated with Accutase (Gibco). Cells (5 × 10^5^) were resuspended in FACS buffer (D-PBS containing 2% FBS) with APC-CD44 (1:25; BD Pharmingen) and CD24-PE (1:50; BD Pharmingen) conjugated antibodies and incubated at 4°C for 30 min. Cells were washed with cold FACS buffer, resuspended in cold FACS buffer, and subjected to flow cytometry. Data were plotted as CD24-PE versus CD44-APC to identify CD44^hi^/CD24^lo^ populations based on single stained controls.

### Tumorsphere Assays

Single cell suspension was filtered through a 40-µm sieve (BD Falcon) and seeded in ULA plates (2000 cells per well). Cells were grown in supplemented phenol-red free DMEM/F12 as described above (3D culture conditions). For secondary tumorspheres, primary tumorspheres were dissociated in 0.25% trypsin-EDTA. Cells were plated as described above in supplemented phenol-red free DMEM/F12. Tumorspheres were allowed to form for 7-10 days. Tumorspheres were analyzed by total number and scored by manual counting using a scaled grid. Data are presented as the average ± SD of three independent measurements.

### Co-Immunoprecipitation Assays

Cells were harvested in ELB lysis buffer [50 mM HEPES, 0.1% NP-40, 250 mM NaCl, 5 mM EDTA, 1X complete protease inhibitors (Roche), 1X PhosSTOP (Roche), and supplemented with 1 mM PMSF, 1mM NaF, 0.5 mM Na_3_PO_4_, 25 mM BGP, and 20 µg/ml aprotinin]. 1000 µg lysate (1 mg/ml) was incubated with 1 µg of the indicated antibody overnight at 4°C. Immunocomplexes were isolated with protein G agarose (Roche) for 2 h at 4°C. Resin was collected and washed with cold ELB buffer. IgG antibodies were included as controls. Immunocomplexes were eluted with sample buffer, resolved by SDS-PAGE, and analyzed by Western blot.

### Proximity Ligation Assay (PLA)

Cells were plated in 8-well chamber slides and fixed in 4% paraformaldehyde for 15 min at RT. Slides were permeabilized for 15 min in PBS containing 0.1% Triton X-100 and processed using the Duolink Red Mouse/Rabbit Kit (Sigma) per manufacturer’s protocol (Sigma). Slides were incubated with primary antibody (1:250 to 1:1000) overnight at 4°C. PLA was performed with the following antibodies: ERα (F-10, Santa Cruz Biotechnology), PR (H-190, Santa Cruz Biotechnology). The ratio of puncta/nuclei for each experimental condition was calculated by counting all puncta and nuclei in 40x images. Confocal microscopy was used to image slides.

### Soft Agar Assays (Anchorage Independent Growth)

Cells were seeded (4 × 10^4^ cells/well) in 1X sterile low melt agarose (Invitrogen) containing 5% DCC and the appropriate treatment (vehicle [EtOH], E2 [1 nM]). Soft agar assays were allowed to proceed for 18-21 days at 37 °C. Afterwards, cell colonies were counted with ImageJ (Wayne Rasband, National Institutes of Health, http://rsbweb.nih.gov/ij/). Data are presented as the average ± SD of three independent measurements.

### Mammary Intraductal (MIND) Model

Intraductal injections were performed as described (34) in seven-week old female NOD.Cg-*Prkdc^scid^ Il2rg^tm1Wjl^*/SzJ (NSG) mice (#005557, Jackson Laboratory). Mice were anesthetized with 2% inhaled isoflurane prior to intraductal injections. Five mice/group were injected with 2.5 × 10^4^ cells into each nipple of the 4^th^ inguinal glands with the indicated breast cancer cell line. Engrafted mammary glands were harvested 8 weeks after injection, fixed in 4% paraformaldehyde, and processed for H&E staining or immunohistochemistry. All animal studies were compliant with ethical regulations approved by the University of Minnesota IACUC committee.

### Histological Analysis

Tissues were fixed in 4% paraformaldehyde and sectioned for hematoxylin and eosin (H&E) staining. H&E sections were analyzed using QuPath to quantitate the total mammary gland area (%) that contained tumor cells. Two-pixel classifiers were used to identify tissue and tumor regions, respectively. Tumor regions were split into individual areas and measured as previously described (51). The total area of the resulting tumor regions was divided by the area of the tissue section times 100 to obtain percent coverage.

### Detection of Circulating Tumor Cells (CTCs)

CTCs were isolated at the end of the MIND xenograft experiment. Fresh blood samples were collected by cardiac puncture and processed by density centrifugation using Isolymph/Ficoll-Paque (Sigma) as described previously (52). Buffy coat containing CTCs was seeded into DMEM containing 5% FBS and 1% penicillin-streptomycin, and 1X sterile low melt agarose (Thermo Fisher). Colonies were grown for 14 days at 37 °C. Data are presented as the average ± SD of five independent measurements.

### RNA-Sequencing

#### Sample collection and sequencing

For 2D conditions, T47D WT and Y537S ESR1 cells (shGFP and shPR) were plated in triplicate. Cells were hormone-starved for 24h, followed by treatment with E2 (1 nM, 48h) or R5020 (10 nM, 6h). For 3D conditions, cells were plated in triplicate into tumorsphere media in ULA plates and allowed to form for 6 days. RNA extraction was performed using RNeasy Mini Kit (Qiagen). Purity of the total RNA samples was assessed using BioAnalyzer (Agilent) and samples with an RNA integrity score >8 were used for library construction by the University of Minnesota Genomics Core. Libraries were built from individual samples with the TruSeq Stranded mRNA kit (Illumina). 50 base pair paired-end sequencing was performed using Illumina NovaSeq X Plus. On average >15 million reads were sequenced per sample.

#### Data analysis

Quality scores across sequenced reads were assessed using FASTQC. Illumina adapters were removed using Trim-Galore. For alignment and transcript assembly, the sequencing reads were mapped to hg38 using STAR (53). Sorted reads were counted using HTseq (54) and differential expression analysis was performed using DESeq2 (55). Genes with a p-value of <0.05 and a log2 fold change greater than 1 or less than −1 were considered differentially expressed. Comparative gene set enrichment analysis (GSEA) was performed using the ClusterProfiler package (56) to identify differentially enriched pathways between experimental conditions.

### ChIP-seq

#### Sample collection and sequencing

T47D WT and Y537S ESR1 cells (shGFP and shPR) were hormone-starved for 48h and then treated with vehicle or R5020 (10 nM, 1h). 1 × 10e^6^ cells were harvested in ice-cold PBS. Cells were crosslinked in 1% formaldehyde in PBS. Crosslinking was quenched by adding glycine at a final concentration of 125 mM. Crosslinked cell pellets were snap-frozen and stored at −80°C. For each ChIP experimental replicate, 2 × 10e^6^ crosslinked cells (from 2 crosslinked aliquots) were lysed in lysis buffer with PICS III using sonication (high, 30 seconds on/off, for 5 intervals of 10 minutes). 5% of lysate was reserved for input control and snap frozen to store at −80°C. Lysates were diluted to 1 µg/µl protein based on Nanodrop A280 concentrations and divided into 1 ml aliquots. 5 µg of the appropriate antibodies (phospho-PR [S294] for phospho-PR ChIP, rabbit IgG for phospho-PR negative control) were added to the appropriate lysate aliquots and rotated at 4°C overnight. Protein-chromatin was isolated and eluted using protein G beads (Cytiva Streptavidin Mag Sepharose Magnetic Beads, Thermo Fisher). Eluted ChIP samples were incubated with RNAse A and Proteinase K to reverse the crosslinked protein-chromatin. Input samples and ChIP-DNA were purified using a Qiagen QIAquick PCR Purification Kit, and purified DNA samples were eluted in 30 µl nuclease-free water. ChIP-DNA was quantitated by Qubit (Life Technologies), libraries were generated using the Illumina ChIP-SEQ kit and quality assessed using Bioanalyzer (Agilent). Sequencing was performed on the Illumina NovaSeq X by the University of Chicago Functional Genomics core. Samples were analyzed over 50M clusters per sample, which was variable due to the available library content.

#### Data analysis

Initial quality control and peak calling was performed using Nf-core ChIPseq pipeline (57). DeepTools v.3.5.6 (58) bamCompare function was used to generate fold enrichment bigWig tracks for each sample compared to the IgG control and to make enrichment plots. DiffBind (59) was used to identify regions of differential accessibility. HOMER (60) was used for transcription factor motif analysis and peak annotation. GEO accession: XXX

### Statistical Analysis

Analyses were performed using GraphPad Prism. Data were tested for normal distribution using Shapiro-Wilks normality test and homogeneity of variances using Bartlett’s Test. Once data met these two requirements, statistical analyses were performed using one-way or two-way ANOVA in conjunction with Tukey multiple comparison test for means between more than two groups or Student *t* test for means between two groups, where significance was determined with 95% confidence.

### Data Availability Statement

GEO accession numbers will be provided.

