## Supplemental Material for "Progesterone receptors drive advanced breast cancer phenotypes including circulating tumor-and stem-like cell expansion in the context of *ESR1* mutation"

**SUPPLEMENTARY MATERIAL**

**SUPPLEMENTARY FIGURES AND LEGENDS**

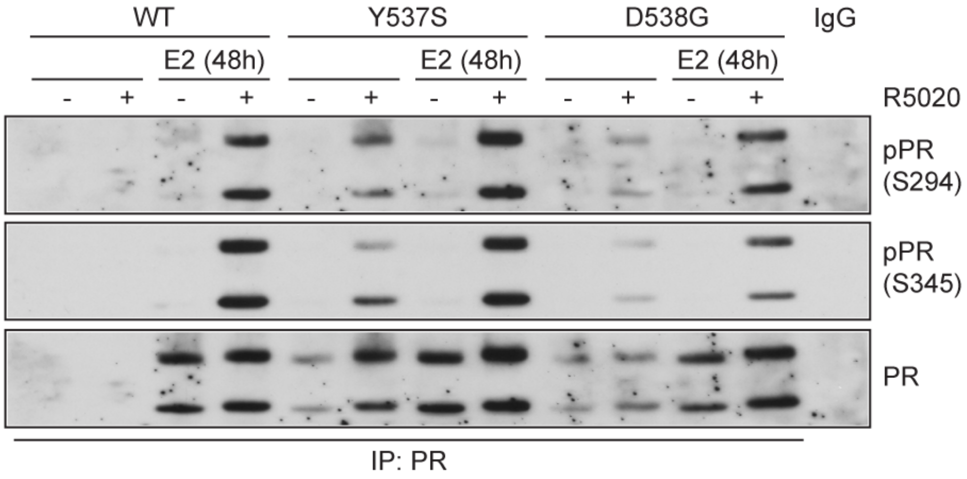


**Supplementary Figure 1.** PR phosphorylation levels in MCF7 ER cells. Immunoprecipitation of PR in MCF7 ER (WT, Y537S, D538G) cells pre-treated with E2 (1 nM) for 48 h followed by treatment with veh (EtOH) or R5020 (10 nM) for 60 min.

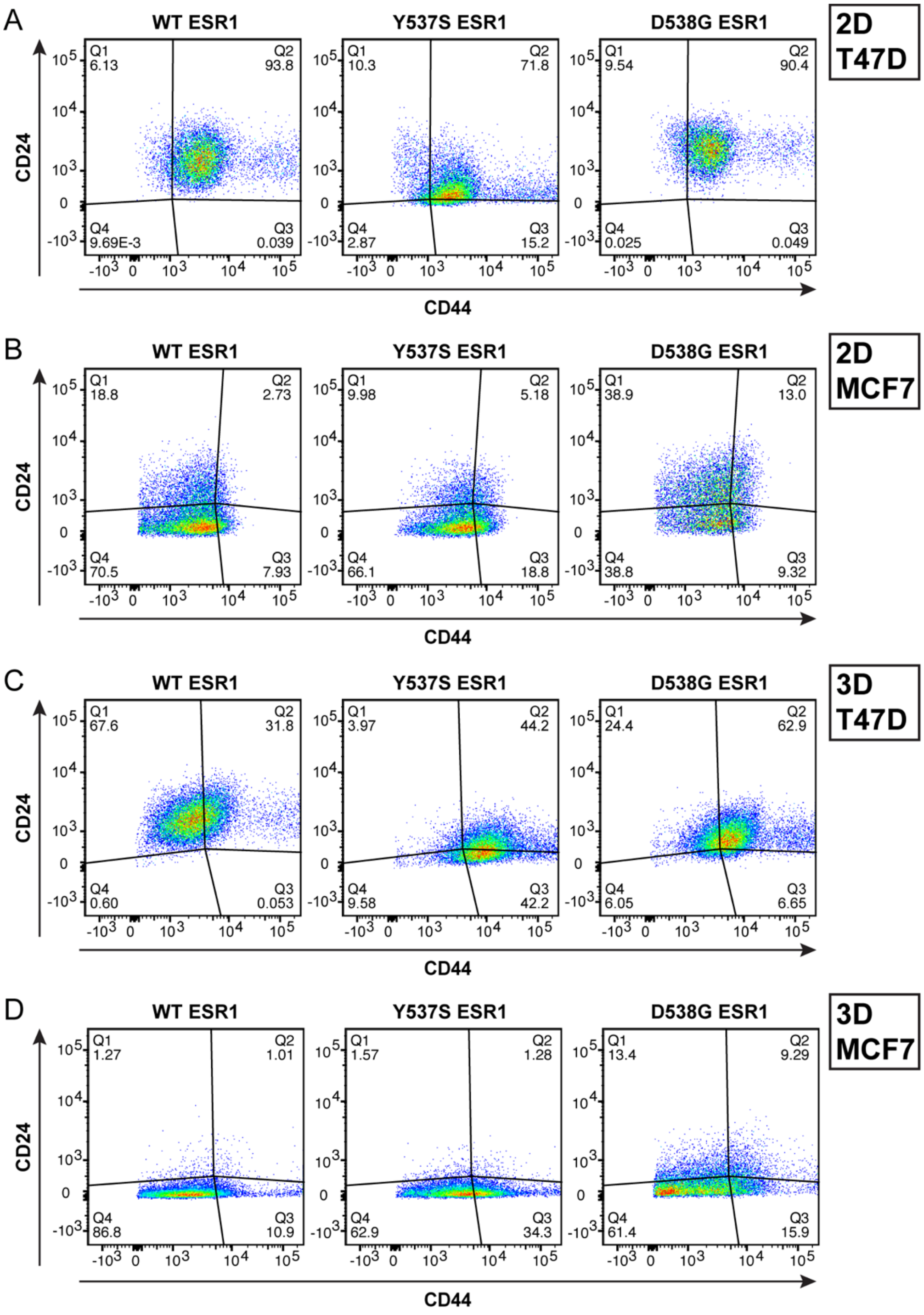


**Supplementary Figure 2.** Representative flow cytometry dot plots shown for CD44$^{hi}$/CD24$^{lo}$ populations in T47D and MCF7 ER cells cultured in 2D (adherent) (**A, B**) and 3D (tumorsphere) (**C, D**) conditions.

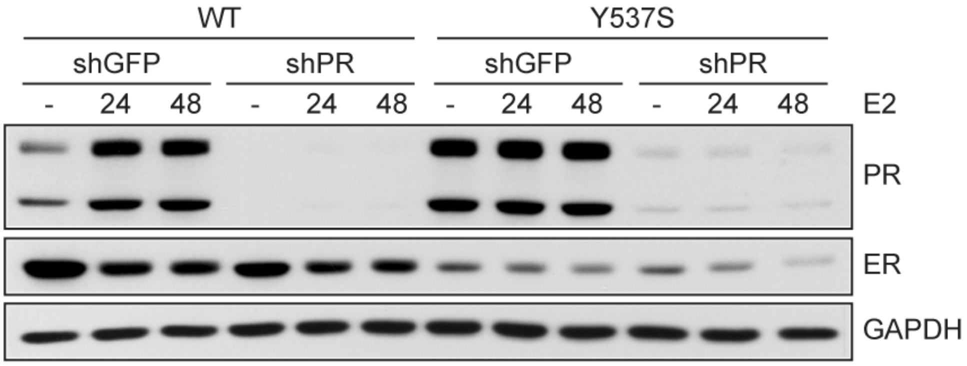


**Supplementary Figure 3.** Western blot of PR knockdown in T47D ER (WT, Y537S) cells treated with E2 (1 nM) for 24 and 48 h.

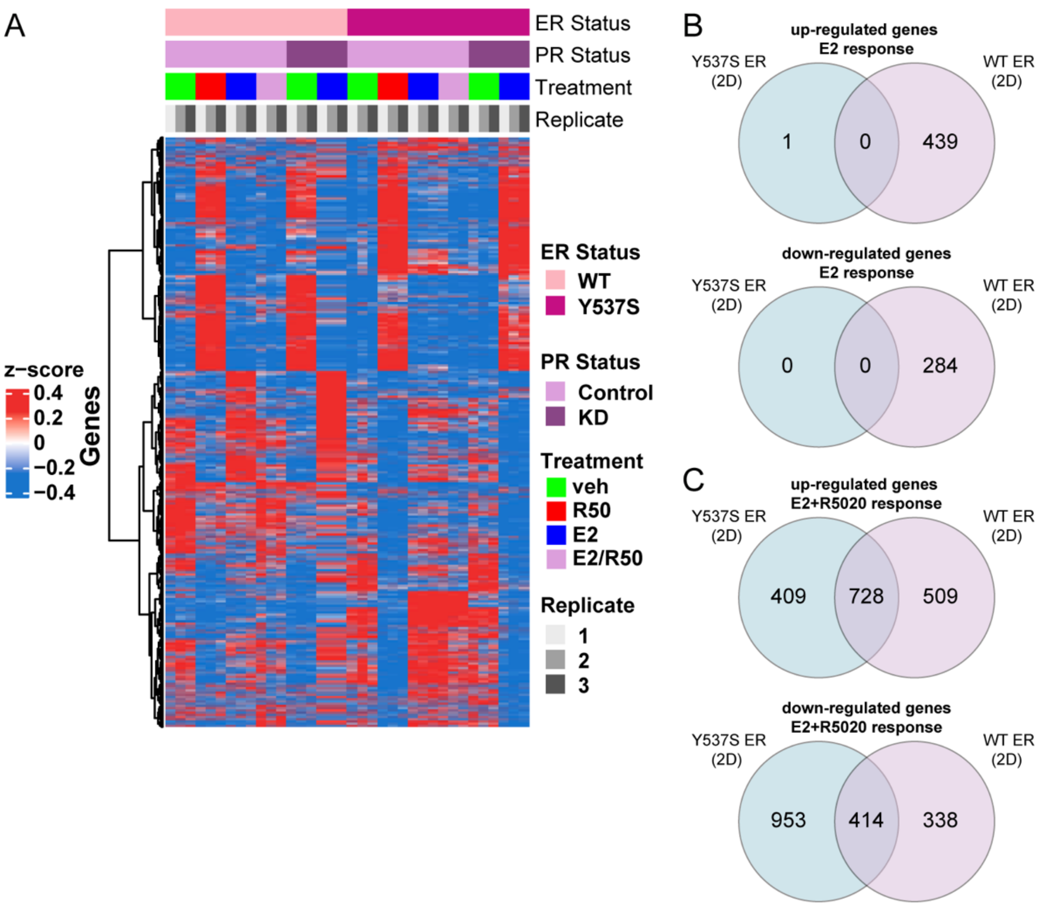


**Supplementary Figure 4.** (**A**) Hierarchal clustered heat map of 2D samples. Venn diagrams showing differentially expressed genes up or downregulated >4-fold in response to (**B**) E2 (1 nM) or (**C**) E2 (1 nM) plus R5020 (10 nM) combination.

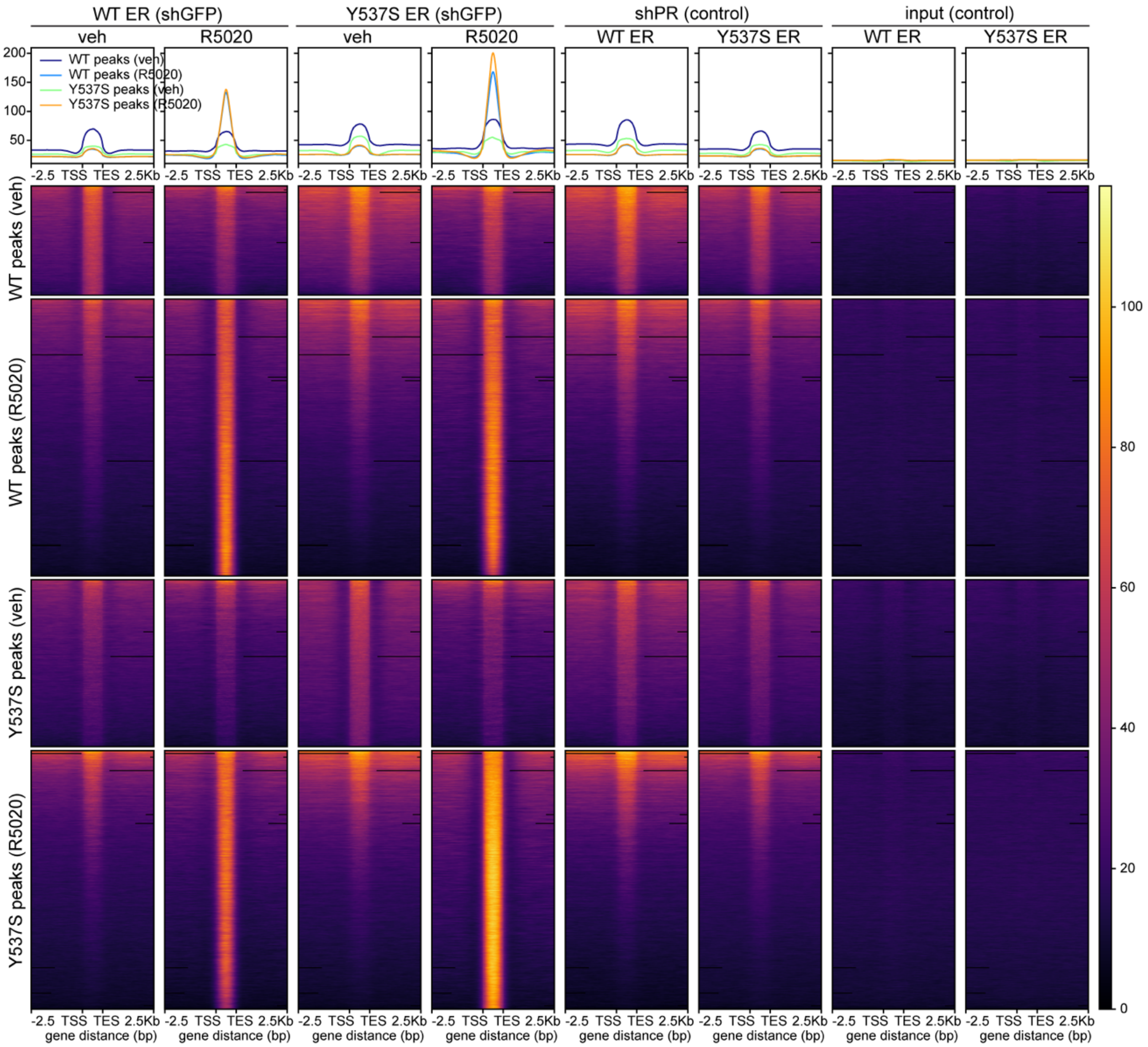


**Supplementary Figure 5.** Phospho-PR ChIP-seq signal in WT and Y537S ER cells in response to R5020 (10 nM). shPR and input samples are included to show enrichment compared to negative controls.

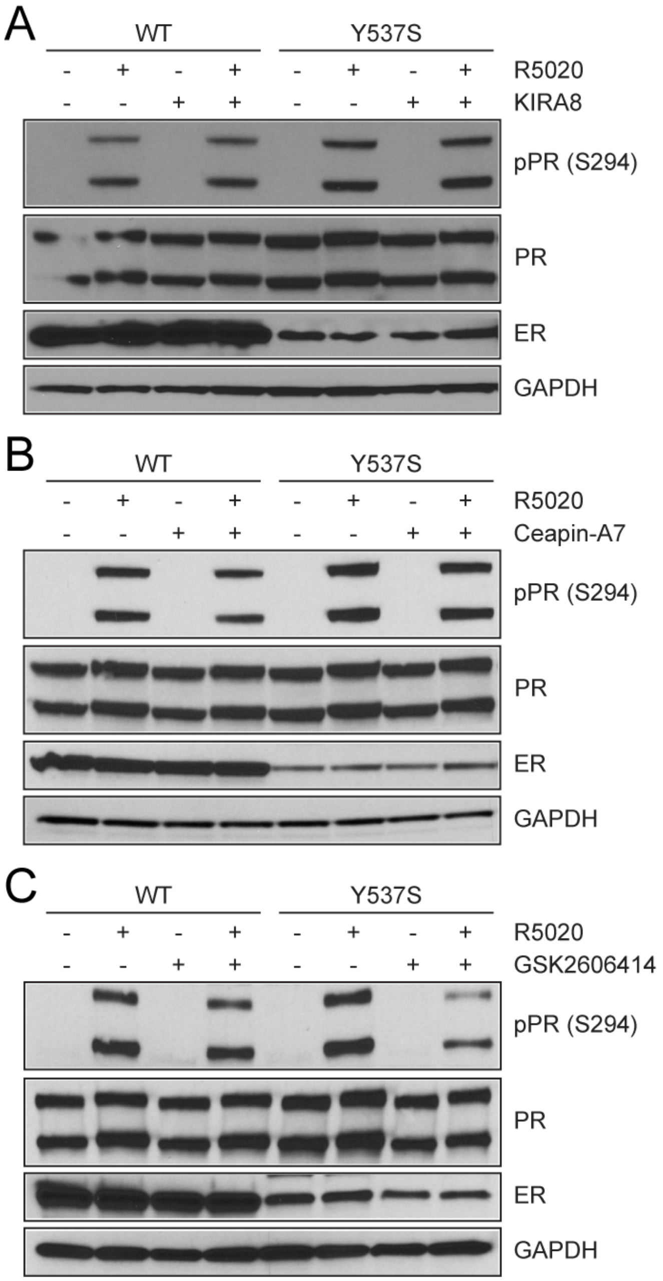


**Supplementary Figure 6.** Western blot of T47D ER cells (WT, Y537S) pre-treated with vehicle, (**A**) KIRA8 (5 µM), (**B**) Ceapin-A7 (5 µM), or (**C**) GSK2606414 (5 µM) for 2h followed by vehicle or R5020 (10 nM) for 1h.
